# Facial Expressions of Emotion are Infrequent in Toddlers’ and Caregivers’ Egocentric Views. An Ecological Study

**DOI:** 10.64898/2026.02.07.704546

**Authors:** Emma J. Jackson, Elena Geangu

**Affiliations:** Department of Psychology, University of York, York, United Kingdom

**Keywords:** Toddlers, Emotion, Egocentric Vision, Sparsity, Ecological

## Abstract

Toddlerhood is a critical period in the development of facial expression processing. Prior research suggests that in the natural environment, the frequency of faces in the toddlers’ egocentric view declines relative to infancy. However, the specific statistics of the emotional facial expressions available to the developing toddler remain unknown. This study implemented a dual-perspective set-up to record the egocentric view of toddlers and their caregivers during everyday situations at home (*N* = 26 families). Using automated computer vision models, we quantified both the frequency of faces and the emotional expressions displayed. Confirming our hypotheses, faces were sparse in toddler views and significantly less frequent than in caregiver views. Across both perspectives, happiness was the dominant expression, while negative facial expressions were extremely rare. Notably, faces expressing surprise were frequent in toddler view, whereas caregivers encountered significantly more happy and sad facial displays than their children. This is the first ecological study to objectively quantify the occurrence of emotional facial expressions in the home environment. These findings challenge the assumption of an abundance of emotional signals in the early development. Instead, they demonstrate that toddlers develop face representations based on sparse input that is biased towards positive expressions (e.g., happy), suggesting high efficiency in extracting and generalizing information from limited input.

## 1 Introduction

The human face conveys a rich array of socially relevant visual signals that observers can use to infer others’ intentions, thoughts, and affective states (e.g., Crivelli and Fridlund, 2019; Martin et al., 2017; Jack et al., 2014, 2016). Sensitivity to such facial cues is fundamental for adaptive social functioning and represents a critical developmental task (Denham et al., 2011). Accordingly, a substantial body of developmental research has examined how infants and toddlers come to use facial information to guide their understanding of others’ emotions and behaviour. Collectively, these studies indicate that the abilities to extract and use affect-relevant information have a protracted development spanning the entire childhood and continuing into adolescence (e.g., Herba and Phillips, 2004; Meltzoff and Ryczek, 2025; LoBue et al., 2025; Ruba and Repacholi, 2020; Riddell et al., 2024).

Behavioural studies using looking-time paradigms found that during the first months of life, infants gradually start to perceptually discriminate between facial displays of happiness, fear, sadness, and anger (e.g., Farroni et al., 2007; Field et al., 1983; Young-Browne et al., 1977). By 5–7 months, they also appear to detect transitions between different facial expressions (Kotsoni et al., 2001). Around the same age, infants begin grouping faces based on discrete emotion expressions; initially, this seems to be predominantly the case for happy facial expressions when contrasted with anger and fear (e.g., Geangu et al., 2016a; Ruba and Repacholi, 2020; Safar and Moulson, 2017). Evidence for categorical grouping for negative facial expressions, particularly within valence, tends to be found in older infants (e.g., Ruba and Repacholi, 2020), although this ability seems to depend on the task. Indeed, when the complexity of the visual environment increases to include multiple categories, Woodard et al. (2022) found that only children 5 years and older differentiate based on discrete emotion categories, while younger 3- to 4-year-olds predominantly discriminate based on valence.

Research into the development of infants’ own expressivity provides further clues regarding when they begin to process the communicative value of emotional facial expressions. Between 2 and 6 months, infants spend increasing amounts of time smiling (Malatesta et al., 1986, 1989), and the display of different smile configurations during face-to-face interactions becomes heavily linked to their mother’s facial signals. During this period, infants become increasingly likely to use smiling configurations that indicate high arousal (i.e., open-mouth smiling with eye constriction) when they are looking at their mother while she is smiling and looking at them (Messinger and Fogel, 2007). This suggests that during the first months after birth, infants learn the potential communicative value of smiling faces during face-to-face interactions.

Toward the end of the first year of life and during toddlerhood, there are indications that children use emotional facial configurations to guide their behaviour in different contexts. Several studies have shown that during novel situations (e.g., encountering novel toys), infants are more likely to show avoidant behaviours if their mothers show negative facial expressions towards the novel event, and are more likely to approach if the mothers express positive affect (e.g., Gerull and Rapee, 2002; Mumme and Fernald, 2003; Sorce et al., 1985; Tamis-LeMonda et al., 2008; Walle et al., 2017). Later, in the second year of life, children appear to embed facial expressions within the context of people’s actions, which modulates how expressions affect their behaviour. For example, 19-month-old (but not 16-month-old) infants respond with congruent negative affect to caregiver painful expressions, but only when these are preceded by a genuine accident. By 24 months, toddlers begin to show differentiated behavioural responses to a variety of others’ discrete emotions (e.g., Özden et al., 2025; Walle et al., 2017) and associate prototypical facial expressions with a variety of emotionally relevant events (e.g., Denham, 2019). Toddlerhood also represents the age when children begin to associate facial expressions of emotions with corresponding verbal labels. Importantly, however, like many aspects of affect knowledge, this ability continues to improve throughout childhood (Riddell et al., 2024).

Taken together, these findings suggest that toddlerhood represents a period of continuing dynamic development in children’s representations of facial emotion expressions. But what drives these developmental changes? While the points of inflection in developmental outcomes have been well documented, less is known about the mechanisms and processes that drive these changes. Although theoretical proposals have been formulated in this respect (e.g., Hoemann et al., 2020; Leppänen and Nelson, 2009), the field is still far from providing compelling explanations. One promising avenue is to consider emotion understanding and facial expression processing abilities as outcomes of a cascade of developmental events (e.g., Hoemann et al., 2020; Masten and Cicchetti, 2010; Oakes, 2017). According to this view, the milestones illustrated above reflect specific experiences that have shaped relevant physical, motor, affective, and cognitive abilities at various points during the developmental timeline. This is particularly relevant because in the natural environment, emotions are multidimensional, dynamic, and inherently interactive events (e.g., Reschke et al., 2018). Infants are not mere spectators; they are active perceivers engaged in social transactions where others’ expressive signals are embedded within the context of their own emotional responses and the physical environment (Geangu, 2015; Özden et al., 2025; Smith et al., 2018; Davidov et al., 2025). Changes in the child’s own motor, cognitive, and emotional abilities lead to changes in how they perceive and understand people’s behaviour, and how they interact with people and their environment (e.g., Csibra and Gergely, 2013; Denham et al., 2011; Geangu et al., 2015; Phillips et al., 2002; Poulin-Dubois et al., 2018; Reschke et al., 2017a; Teufel et al., 2010). These alter the quality and quantity of their experiences, which in turn drives further developmental change (e.g., LoBue et al., 2025; Oakes, 2017; Smith et al., 2018).

One key contribution of the developmental cascades framework is to bring back into focus the role of input, which resonates with recent calls for developmental psychology to broaden its research portfolio to include objective measures of children’s experiences in everyday life (e.g., de Barbaro and Fausey, 2022; Rogoff, 1998, 1997; Rogoff et al., 2018; Warlaumont et al., 2022). The idea that input is relevant for development, including the development of face processing, is not new (e.g., Bruner, 1983; Gibson, 1982, 1988; Johnson, 2011; Leppänen and Nelson, 2009; Morton and Johnson, 1991). However, the majority of research on face processing and emotion expression recognition remains over-reliant on laboratory-based approaches with simplified stimuli that depict exaggerated and posed facial expressions. These are often chosen based on assumptions about the nature of the input in everyday life, yet the input is rarely, if ever, objectively measured. It is recognized, however, that ‘in the wild’ facial displays are subtle, fleeting, varied, modulated by social norms, and heavily contextualized (e.g., Crivelli and Fridlund, 2019; Denham et al., 2004; Jack et al., 2014; Rychlowska et al., 2017; Martin et al., 2017). Hence, if we are to understand how the abilities to process and use emotional facial expressions develop, it is crucial to understand the nature of the social input from which infants are actually learning.

In recent years, significant technological advances in infant wearable sensors have greatly facilitated such ecological approaches (e.g., Geangu et al., 2023; Long et al., 2022a; Smith et al., 2018, 2015), showing immense potential for seminal discoveries. Head-mounted cameras worn by infants and toddlers in their natural home environment have revealed that their egocentric views are characterized by a non-uniform occurrence of faces (e.g., Jayaraman and Smith, 2019; Fausey et al., 2016; Long et al., 2022a; Sugden and Moulson, 2019; Smith et al., 2015). During the first months of life, faces occur with the highest frequency in infants’ views (approximately 25%), but this exposure decreases substantially in the following months; by 24 months, faces appear in the toddler’s view for only about 7.5% of the time (Fausey et al., 2016; Jayaraman et al., 2015, 2017; Long et al., 2022a; Sugden and Moulson, 2019), before increasing by adulthood to rates similar to those of young infants (e.g., 23%, Oruc et al., 2019). During the early months, faces also tend to be visually larger, more proximal, and temporally dense (Jayaraman and Smith, 2019; Long et al., 2022a). Importantly, caregivers’ faces are most prevalent, appearing consistently across physical contexts (Long et al., 2022a; Sugden and Moulson, 2019). While face presence declines with age, the infant and toddler view becomes increasingly dominated by hands and objects (Jayaraman and Smith, 2019; Long et al., 2022a). This shift may be partially driven by changes in locomotor ability (e.g., Karasik et al., 2014; Kretch et al., 2022, 2023). Unlike younger infants, crawling and toddling infants often must actively look up to see adult faces. This shift may also be the cascading effect of changes in cognitive abilities, as infants become increasingly interested in exploring and learning about the physical environment (e.g., Fausey et al., 2016; Smith et al., 2018).

However, despite these substantial advances in characterizing the regularities with which infants and toddlers are exposed to facial information in their natural environment, to our knowledge there are no studies that objectively measure and document the extent of young children’s exposure to emotional facial expressions. The necessity of characterizing this input is underscored by indirect evidence suggesting that variations in exposure, whether experimental, cultural, or clinical, shape processing strategies. Some of this evidence comes from studies showing that already within a few days of birth, infants seem able to learn from the visual input available to them. For example, Addabbo et al. (2018) showed that while newborns do not spontaneously discriminate between dynamic facial expressions of happiness and disgust, if they are given the opportunity to become familiarized with these expressions through repeated presentations, they show a pattern of visual preference indicating perceptual discrimination.

Other streams of research have suggested that cultural variations in the facial expressions infants are exposed to may shape perceptual strategies for sampling relevant visual information. Using data-driven approaches for modeling fixation patterns, Geangu et al. (2016a) showed that Western-Caucasian 7-month-old infants (born and raised in the UK) manifest a visual scanning pattern distributed across the entire face, although they look more at the mouth area of fearful and happy faces, while East-Asian infants (born and raised in Japan) focus on the eyes. These culturally specific information sampling strategies closely resemble those previously reported for adults in these cultures (Jack et al., 2012). Importantly, while strikingly different, both strategies are effective for discriminating between facial expressions. It has been hypothesized that these visual scanning strategies may be partly driven by cultural variations in input. Japanese mothers value expressing less affect in front of their children, and have been reported to use less emotional facial expressivity and more non-direct body contact during interactions with their infants compared to Western mothers (Denham et al., 2004; Fogel et al., 1988). These findings resonate with a large body of work on cultural variations in affect display rules, showing that Japanese adults value reduced facial expressivity (Safdar et al., 2009), which may lead to the increased attention to culturally specific facial signals in the eye region.

Research involving clinically at-risk populations provides further supporting evidence that the quantity and quality of input may be relevant for the development of emotion expression processing. For example, infants raised by depressed mothers tend to show different responses to emotional facial configurations during lab-based experiments (e.g., less interest in negative expressions, prolonged visual attention to happy facial expressions) (Cohn et al., 1986; Diego et al., 2004; Field, 1992) compared to infants raised by non-depressed mothers. This has been conjectured to result from increased exposure to negative affect, since depressed mothers tend to express more negative affect compared to non-depressed mothers (Aktar et al., 2017; Murray et al., 1993). Other clinical studies have shown that relative to children from typical families, children raised in families characterized by physical aggression show enhanced abilities to recognize and allocate more attention to angry faces compared to other facial expressions of emotions (Cicchetti and Curtis, 2005; Curtis and Cicchetti, 2011). Collectively, these findings suggest that the developmental system is highly sensitive to the statistical regularities of the emotional environment, developing processing strategies for the predominant features of the input.

Given that the actual characteristics of the input supporting the development of rich emotional representations during toddlerhood remain largely unknown, the present study aims to objectively quantify these visual regularities in the natural environment. To achieve this, we employed a dual-perspective set-up to record the egocentric views of both toddlers and their caregivers in the natural home environment (Figure 1). This simultaneous recording approach captures the dynamic, interactive context in which emotional facial expressions occur. To ensure the ecological validity of the data and capture a diversity of emotion-eliciting situations, we recorded several hours of real-life experience sampled at different times within a week. Finally, to facilitate the analyses of this extensive dataset, we developed a specialized pipeline for automated face detection and emotion expression estimation, leveraging recent advances in computer vision and deep learning.

**Figure 1:**
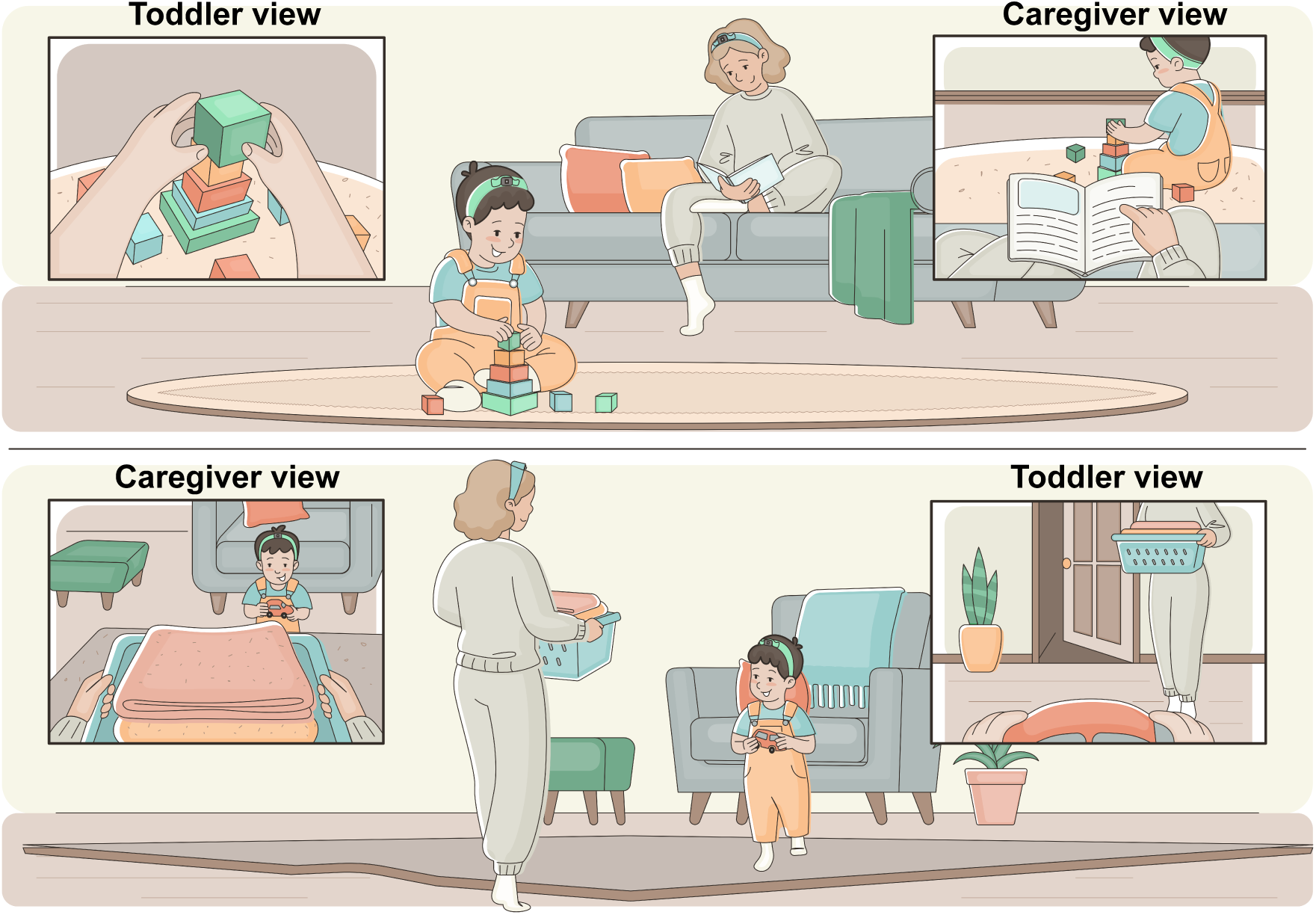
Illustration of a dual-perspective recording set-up in the home environment using HMCs worn by a caregiver and a toddler. Two everyday scenarios are depicted. Insets show illustrative egocentric views, based on head orientation and activity context. These views are not intended as precise depictions of the human and HMC fields of view, but rather as conceptual representations of what is likely to be in the views of the toddlers and caregivers during naturalistic interactions.

In the absence of prior quantitative data on emotional facial expressions input, we inform our predictions from the functional framework of parent-child emotion regulation. Developmental research characterizes toddlerhood as a period of heightened emotionality and immature self-regulation, requiring caregivers to act as ‘external regulators’ who closely monitor the child’s affective states (e.g., Buss and Kiel, 2004; Cole et al., 2004; Denham, 2019). This dynamic creates a distinct functional asymmetry between the two partners. On one hand, toddlers are likely to experience and express intense emotions. During episodes of play, distress or frustration, the caregiver’s attention is also more likely to be fixated on the child to assess needs and ensure safety. On the other hand, caregivers are more likely to regulate their emotions and facial expressivity of negative affect to modulate children’s arousal (Denham et al., 2004). Based on these functional roles, we formulate two key predictions. First, regarding face presence, we predict that faces will occur less frequently in toddlers’ compared to caregivers’ views, replicating previous findings. Second, regarding the emotional content, we predict that the toddlers’ view will be characterized by a lower frequency of emotional displays than the caregivers’ view, and specifically a marked scarcity of negative affect.

## 2 Methods

### 2.1 Participants

Forty-one toddler-caregiver pairs (22 female and 19 male toddlers) were invited to participate in the study. They were recruited from in and around an urban area in the United Kingdom (UK). Nine toddler-caregiver pairs did not return any data, 6 pairs returned data deemed of insufficient quality for analyses, and 26 toddlers provided analysable data. Our final sample size is comparable to or exceeds those reported in previous studies per age group (e.g., Fausey et al., 2016; Jayaraman et al., 2015, 2017; Jayaraman and Smith, 2019; Long et al., 2022b,a; Sugden et al., 2014). Detailed information about inclusion criteria is provided below. On average, toddlers included in the analyses were 29.26 months old (female *M_age_* = 29.37 months, *SD_age_* = 0.56 months; males *M_age_*= 29.08 months, *SD_age_* = 0.62 months). Caregivers provided informed consent before the beginning of the study procedure. The study was approved by the Psychology Department Research Ethics Committee.

### 2.2 Technical equipment

Egocentric video was captured using lightweight mini cameras (Magendara, 23g, Figure 2A, B). The cameras recorded video at 1280*×*720 resolution (15 fps), with a diagonal field of view (FOV) of approximately 70*^◦^*, corresponding to a geometric vertical FOV of approximately 40*^◦^*. To optimize the viewing range for capturing faces, the main objective of our study, the cameras were attached to headbands and placed approximately 8cm above the nasion for toddlers (Figure 2D). This placement likely biases the camera upwards relative to the toddler’s natural line of sight. Previous studies suggest that faces tend to be concentrated in the middle-upper part of the toddler’s view captured by head-mounted cameras (e.g., Fausey et al., 2016; Long et al., 2022a,b). Furthermore, in a previous validation study using head-mounted eye-tracking (Zhang et al., 2025), it was observed that toddler fixations during naturalistic scenarios were highly concentrated within the central visual field, with a marked decrease in fixation probability in the extreme upper periphery. This is consistent with a wider literature indicating a ‘centering bias’ (e.g., Bambach et al., 2016a,b, 2017; Franchak et al., 2021, 2024; Borjon et al., 2021), where head movements are actively recruited to bring objects into the center of the field of view for processing, as well as research showing that adults rotate both head and eyes to facilitate visual exploration in different contexts (e.g., Franchak et al., 2021).

**Figure 2:**
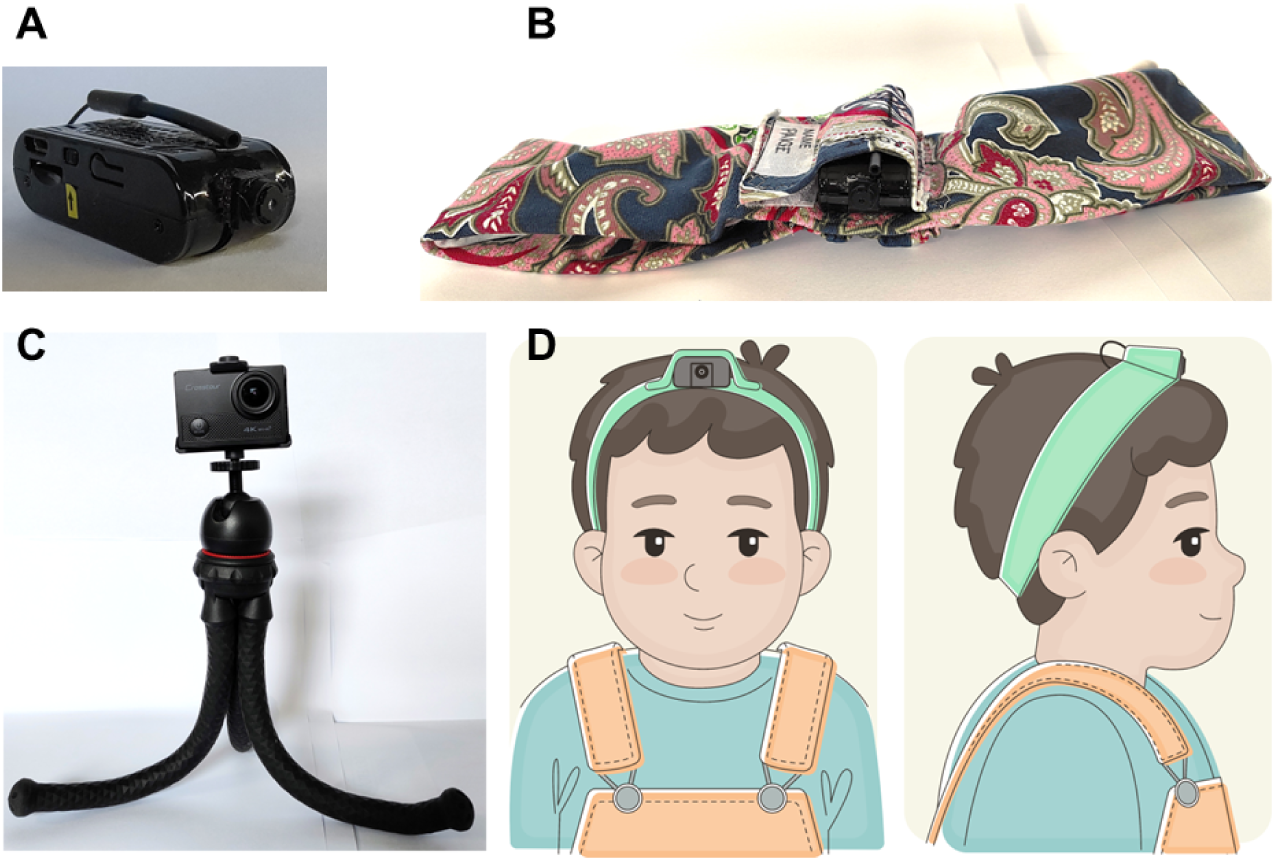
The equipment used in the study: A) the head-mounted camera; B) the head-band with the integrated head-mounted camera; C) the room camera mounted on a flexible tripod; D) a schematic illustration of the head-mounted camera worn by a toddler.

Each family received a recording kit containing two head-mounted cameras to record the egocentric perspectives (one for the toddler and one for the caregiver); one stationary room camera with a tripod to record the wider context of the interaction environment and verify the head-mounted camera data validity; and instruction sheets (Figure 2).

For both toddler and caregiver, the head-mounted camera was attached to a breathable head-band, designed for comfort and maintaining the best position for capturing the wearers’ egocentric view during free movement (Figure 2 B). To adapt to the head circumference of the toddlers and the caregivers, two separate headbands were designed. The stationary room camera was mounted on a flexible tripod to enable placement on different surfaces in participants’ homes. This camera was only used to validate the HMC recordings, and the resulting footage was not analysed in this paper.

### 2.3 Procedure

Recordings took place in participants’ homes from January to December 2021. The COVID-19 pandemic necessitated remote procedures and minimal physical contact with families. Families received detailed instruction packs and communicated with the research team via video calls.

For a week prior to data recording, caregivers were asked to fill in a rough daily diary of what their toddler was doing and when over the seven days. Based on this information, researchers suggested the time when recordings should take place for each of the 7 days of data collection. These times were chosen to capture a wide diversity of everyday routines and activities. However, caregivers were allowed to adjust these suggestions based on how their week unfolded, which was often difficult to anticipate.

Each family was informed that the aim of the study was to understand how the typical natural environment is related to toddlers’ social development, without any particular emphasis on which aspects of the environment will be measured. They were asked to record one continuous hour of video per day for seven consecutive days, during typical daily activities. This dense sampling schedule was intended to facilitate recording a wide range of experiences and emotion-eliciting events. Caregivers were instructed not to change what they would typically do at that time of the day for the purpose of the recording, but rather carry on with their usual activities as they would normally do.

After receiving the recording kits, and prior to commencing the recordings, researchers met online with the caregivers to instruct them how to operate the devices. Emphasis was placed on strategies for best positioning of the head-mounted cameras to best capture the information appearing in the toddler’s and caregiver’s views (Figure 2). Toddlers and caregivers wore their respective cameras simultaneously during recording sessions while the stationery room camera recorded a third-person perspective.

### 2.4 Video data pre-processing and quality control

To ensure the HMC recordings are representative for wearer’s view, each continuous recording session was watched by researchers in its entirety with the aim of identifying portions where the HMC was not well positioned on the head and/or contained significant image quality deterioration (e.g., blur, occlusion by hair, etc.). Where necessary, the room camera view and/or the other HMC view were used to identify these cases. We implemented two complementary criteria for screening sessions. To filter out very brief recordings where participants did not continue with the session or were not compliant with wearing the camera, we excluded all recordings with less than 10 minutes of valid footage (e.g., the camera was only worn on the head for a few minutes, after which was left recording on a table). To filter out invalid footage due to other quality criteria (e.g., blurred image, view occluded by hair or other clothing), we excluded recording sessions with more than 80% of the duration deemed of insufficient quality. For all sessions retained in the analyses, sections of insufficient quality were manually annotated, and further excluded from the analyses in the final stages of data processing (Figure 3). Only participants with at least one valid recording session of more than 45 minutes were included in the analyses.

**Figure 3:**
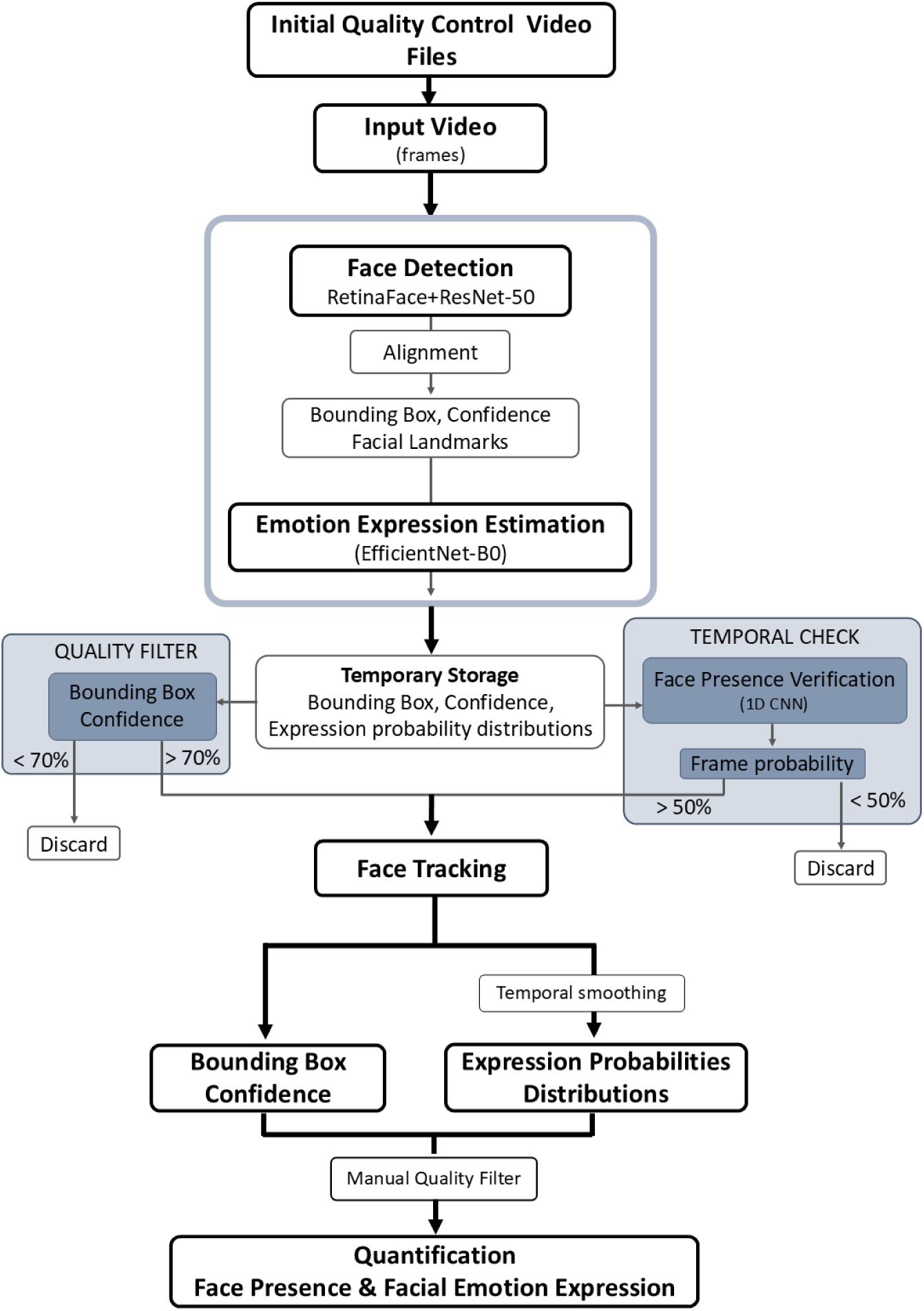
Illustration of the automated face processing pipeline.

From the 32 families who captured any video footage with the HMCs, 26 provided valid data for the toddler-perspective, and 25 for the caregiver perspective. Five families failed to capture any data from the toddler perspective, and one family provided recordings from the toddler’s perspective, but the entire footage was heavily blurry due to a HMC technical failure. Out of the 26 families with usable toddler-perspective data, 24 toddlers had matching caregiver valid recording sessions. Across the entire sample providing valid data, a total of 144 sessions (*M* = 5.54, *SD* = 2.06) of valid toddler-perspective recordings, and a total of 143 sessions (*M* = 5.72, *SD* = 2.03) of valid caregiver-perspective recordings were included in the analyses. On average, the toddler sessions contained 53.42 minutes of valid data (*SD* = 12.17), while the caregiver sessions contained 57.11 minutes (*SD* = 11.42). In total, 6,738,080 frames of toddler footage (*M* = 4.8 hours/toddler, *SD* = 1.83), and 7,292,101 frames of adult footage (*M* = 5.4 hours/caregiver, *SD* = 2.08) were included in the analyses. For a subset of the caregiver-perspective recording sessions, the HMCs were worn by fathers (16 sessions) and grandmothers (16 sessions). From the toddler valid HMC recording sessions, a total of 11.66 hours of video were deemed non-analysable and further excluded from analyses, while from the caregiver valid HMC recording sessions, 6.78 hours were excluded.

### 2.5 Automatic face detection and emotion expression estimation pipeline

Due to the large dataset, manual coding of face presence and facial emotion expressions was deemed unfeasible. Therefore, an automatic approach was implemented leveraging recent advances in computer vision and deep learning.

#### 2.5.1 Model architecture and rationale

We developed a multi-stage pipeline (Figure 3) tailored to the unique challenges of naturalistic recordings. The pipeline includes several modules, implemented interdependently. For detailed information on the pipeline implementation see Supplementary Methods.

##### Face detection

We employed RetinaFace with a ResNet-50 backbone (Deng et al., 2020). This architecture was selected for its state-of-the-art performance on the WIDER FACE ‘Hard’ benchmark, making it capable of detecting the partial, off-angle faces typical of toddler views where standard ‘off-the-shelf’ models often fail (Long et al., 2022a). All candidate faces detected in the frame were marked by a bounding box and associated with a confidence level.

##### Face presence verification

Standard face detectors in egocentric footage frequently generate false positives, which can inflate measurements of face presence frequency and introduce noise into emotion expression estimation. To address the challenge of false positives, we incorporated a custom temporal verification model. We trained a lightweight one-dimensional convolutional neural network (1D CNN) using a dataset of frame-level human annotations. This model analyzes the temporal evolution of detection confidence over a 10-frame window to distinguish between the stable signatures of real faces and the transient spikes of background noise. It is optimized to maximize agreement with human coders (Soft Cohen’s Kappa), and serves as a “temporal gatekeeper”, estimating the probability of face presence based on temporal consistency rather than single-frame appearance (see Supplementary Methods for training protocols and architecture details).

##### Face Tracking

Face detections were linked across frames using a tracking-by-detection procedure based on a modified version of constrained video face clustering (Kalogeiton and Zisserman, 2020). By combining identity embedding similarity with temporal and spatial constraints, this method generates face tracks that serve as an intermediate representation, ensuring that the facial expression data is consistently attributed to distinct individuals, rather than aggregated across multiple people in a scene.

##### Facial emotion expression estimation

For expression estimation, we utilized EfficientNet-B0 (Savchenko and Sidorova, 2024; Savchenko, 2022; Savchenko et al., 2022; Tan and Le, 2019). We prioritized this architecture over larger static models because it is fine-tunined on the Acted Facial Expressions in The Wild (AFEW) video dataset. This gives increased robustness to the motion blur and lighting shifts inherent in our footage, without compromising inference accuracy.

#### 2.5.2 Performance evaluation

To validate the reliability of the automated pipeline, we assessed both the face detection and emotion expression estimation components against established benchmarks and internal quality controls.

##### Per-frame face presence estimation

The custom 1D CNN was trained on a large subset of frames from our dataset (*N* = 990, 027 frames; 18.33 hours), sampled from the toddler and adult perspective across six families.

Frames were manually annotated as containing “one or more faces” if at least one internal feature of a face was fully visible, including real individuals in view or photorealistic depictions (e.g., photographs, mirrors). Annotations were conducted by multiple trained researchers, and subsequently reviewed by the first author to ensure consistency. To assess performance, we calculated standard classification metrics on a hold-out test set from the same domain. The 1D CNN demonstrated high alignment with human annotations, achieving a recall of 0.72, precision of 0.76, and an F1 score of 0.74.

##### Facial emotion expression estimation

To validate the performance of the facial emotion expression model, we curated a dataset of 311 naturalistic video clips representative of everyday social interactions involving infants, children and adults. These clips were extracted from a large corpus of 600 hours HMC footage from our group. The footage was collected in the home environments and laboratory play sessions, and in part contains the data included in this study. Crucially, the test set was specifically designed to represent the range of hard cases typical for this type of device. Clips were selected by researchers to represent segments containing a single face displaying one of the six basic expressions (Anger, Disgust, Fear, Happiness, Sadness, Surprise) or an emotionally neutral expression. The final dataset included 251 video clips recorded in the home environment, and 60 in the lab, with a short mean duration (*M* = 1.80 seconds).

We compared the model’s estimations with ratings from 149 näıve adult participants (115 females, 25 males, 9 other) from Western cultural backgrounds. The majority (93%) of these participants were undergraduate students. Due to the large number of video clips, participants were semi-randomly assigned to rate approximately 78-80 clips each, resulting in an average of 37-38 independent ratings per clip. Clips were presented at both full and half speed with the sound muted.

For each video clip, participants reported: 1) The predominant emotion expressed by the face (7-alternative forced choice: anger, disgust, fear, happiness, sadness, surprise, neutral); 2) Their confidence in this choice (1-5 Likert scale, 1 - “Not at all confident”, 5 = “Very confident”); and 3) How intensely was the emotion displayed (1-5 Likert scale, 1 - “Very weakly”, 5 = “Very strongly”, with N/A option for neutral). Although intensity ratings were recorded, they were not used in this study for model evaluation. To establish a robust human ground truth, we computed confidence-weighted labels by aggregating the confidence ratings for each emotion category. This method ensured that the final classification reflected not just the frequency of a label, but also the certainty of the observers. For example, two assignments of “happiness” with a confidence rating of 5 (total score = 10) would outweigh two assignments of “neutral” with a confidence rating of 1 (total score = 2). The category with the highest cumulative weighted score was designated the *Human Winning Expression*, and the category with the second-highest score was defined as the *Human Runner-up*. Consensus for a clip was calculated as the percentage of individual participants whose response matched the Human Winning Expression. The mean consensus across all clips was 69.41% (*SD* = 21.73).

The automated model generated frame-by-frame probability distributions, which were averaged to derive a clip-level prediction. The category with the highest mean probability was designated the Model Winning Expression. The model demonstrated strong convergence with the human perception. For 62.06% of clips, the Model Winning Expression matched the Human Winning Expression (Table 1). When the criterion was relaxed to include the human runner-up category, the model matched human judgment in 78.14% of cases. Only 5.47% (17/311) of clips showed no correspondence between the model’s top two choices and the human top two choices. The agreement varied by category (Table 1). Happiness (78.1%) and Surprise (77.4%) showed the highest correspondence. Negative expressions showed more variance. Specifically, clips labeled Fear by humans showed lower agreement (25.0%) and were sometimes classified by the model as Sadness (33.3% of cases). This divergence likely reflects a variety of factors and is not solely due to model failure. Fear clips elicited the lowest consensus among human raters (54.14%), and in 77.8% of cases where the model missed fear as the winner, it selected fear as the runner-up, suggesting the expressions themselves were subtle and mixed. Furthermore, the human raters assessed the full-resolution video with access to contextual cues (e.g., body posture, physical environment), and had the possibility to review segments in slow motion. In contrast, the model is constrained to isolated, cropped facial regions and lacks access to the broader scene or body posture cues.

**Table 1:** Confusion matrix table showing agreement between human and model winning facial expression categories.

| Human Label | Model Prediction (%) |  |  |  |  |  |  | Total *N* |
| --- | --- | --- | --- | --- | --- | --- | --- | --- |
|  | Anger | Disgust | Fear | Happiness | Neutral | Sadness | Surprise |  |
| Anger | **60.4** | 1.9 | 3.8 | 3.8 | 5.7 | 15.1 | 9.4 | 53 |
| Disgust | 25.0 | **39.3** | 3.6 | 3.6 | 17.9 | 3.6 | 7.1 | 28 |
| Fear | 8.3 | 0.0 | **25.0** | 8.3 | 16.7 | 33.3 | 8.3 | 12 |
| Happiness | 2.7 | 4.1 | 4.1 | **78.1** | 1.4 | 2.7 | 6.9 | 73 |
| Neutral | 16.1 | 0.0 | 0.0 | 3.6 | **41.1** | 16.1 | 23.2 | 56 |
| Sadness | 5.6 | 2.8 | 5.6 | 5.6 | 8.3 | **72.2** | 0.0 | 36 |
| Surprise | 1.9 | 0.0 | 11.3 | 1.9 | 3.8 | 3.8 | **77.4** | 53 |
| *Total* |  |  |  |  |  |  |  | *311* |

*Note.* Values represent row-wise percentages. The ‘Total *N*’ column indicates the number of ground-truth clips per category. Bold values indicate agreement (diagonal).

### 2.6 Quantification of face presence and facial emotion expression estimations

Frames in which the camera was removed or the video quality was insufficient (e.g., it was dark or the lens was blurry) were excluded from all analyses.

#### Face Presence Quantification

From all valid frames for each participant, we calculated the percentage of frames containing faces. A frame was coded as containing faces if the estimated per-frame probability of face presence exceeded 0.5, and there was at least one bounding box with confidence exceeding 0.7.

#### Facial emotion expression quantification

For each detected face, the dominant facial expression was defined as the category with the highest temporally smoothed probability per face track (see Supplementary methods and Figure 3 for details). We calculated two complementary metrics to characterize the visual environment. (1) *Percentage of all detected faces*. To assess the relative makeup of the encountered social signal, we calculated the frequency of each emotion relative to the total number faces detected. For each participant, the count of faces assigned to a specific category was divided by the total number of detected faces. This metric sums to 100% and represents the relative distribution of emotional expressions within the set of faces in view. (2) *Percentage of all face-opportunities*. To capture the absolute prevalence of emotional exposure over time, we calculated the frequency of expressions relative to the total number of ‘face-opportunities’. The numerator was defined as the cumulative count of faces in a category. The denominator was defined as the cumulative count of all valid frames plus an additional count for every concurrent face within a frame beyond the first. This metric ensures that the simultaneous presence of multiple expressive faces in a frame is weighted, rather than being flattened into a single ‘frame’ event. This approach also avoids the ipsative nature of proportional data (where categories sum to 100%), allowing for independent assessment of expression frequency across perspectives. Percentages were calculated separately for each family and recording perspective (toddler, caregiver).

### 2.7 Analytic strategy

To maximize sample size, we analyzed the presence of emotional expressions separately for each perspective, including the subset of participants with valid HMC footage for the toddler (*N* = 26), and the caregiver (*N* = 25), respectively. For within-perspective analyses, we used the *Percentage of all detected faces* metric to compare the relative dominance of expressions within that group’s visual input. For between-perspectives analyses, we used the subset of families with concurrent valid data (*N* = 24). To compare the incidence of emotional expressions between toddlers and caregivers, we used the *Percentage of all face-opportunities* metric. This metric allowed us to compare absolute frequency of emotional events, accounting for differences both in duration of recording and density of faces encountered by each group. For comparing overall incidence of faces, we used the percentage of frames with faces. Statistical analyses were conducted in SPSS 31. To account for the shared home environment, the partner perspective (toddler vs. caregiver) was treated as a within-subject factor. For repeated-measures analyses, violations of sphericity were assessed using Mauchly’s test (Greenhouse–Geisser corrections reported where necessary). For multiple pairwise comparisons, Bonferroni correction was applied. Differences were considered statistically significant at an adjusted *p < .*05.

## 3 Results

### 3.1 Incidence of Faces in the Toddler and Caregiver Views

Across the 24 families contributing concurrent toddler and caregiver perspective data, the automated pipeline processed 6,640,064 (approximately 123 hours) valid frames for the toddler, and 7,171,310 (approximately 133 hours) for the caregiver. On average, faces occurred in significantly fewer frames for the toddler (*M* = 7.67%, *SE* = .92) than for the caregiver (*M* = 13.81%, *SE* = 1.41), *t*(23) = 4.99, *p < .*001. This represents a large effect, Hedges’ *g* = 0.99 (Figure 4).

**Figure 4:**
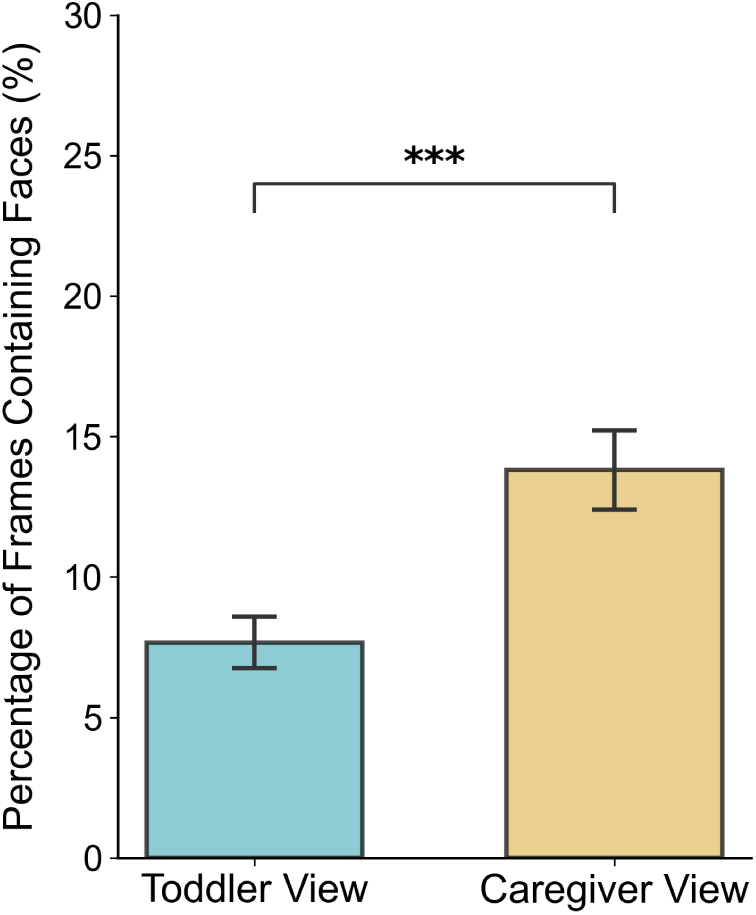
Mean percentage of egocentric video frames containing faces. Data represents the subset of families (*N* = 24) with concurrent recordings. Caregiver views contained significantly more faces on average than toddler views. Error bars represent *±*1 SE. *** *p < .*001.

### 3.2 Presence of Emotional Expressions in the Toddler View

We examined the toddler’s visual input using data from the 26 families with valid toddler-perspective footage. A one-way repeated measures ANOVA showed a significant main effect of facial expression category, *F*(3.21, 80.32) = 207.22, *p < .*001, *η_p_*^2^ = .892. Across all detected faces, *Neutral* faces were by far the most prevalent (*M* = 52.84%, *SD* = 9.34%), followed by *Happiness* (*M* = 13.58%, *SD* = 5.94%) and *Surprise* (*M* = 11.86%, *SD* = 6.14%). Negative expressions were less frequent, with *Sadness* (*M* = 8.34%, *SD* = 4.32%), *Fear* (*M* = 5.10%, *SD* = 3.02%), *Anger* (*M* = 5.01%, *SD* = 5.84%), and *Disgust* (*M* = 3.28%, *SD* = 2.50%) forming the tail of the distribution (Figure 5).

**Figure 5:**
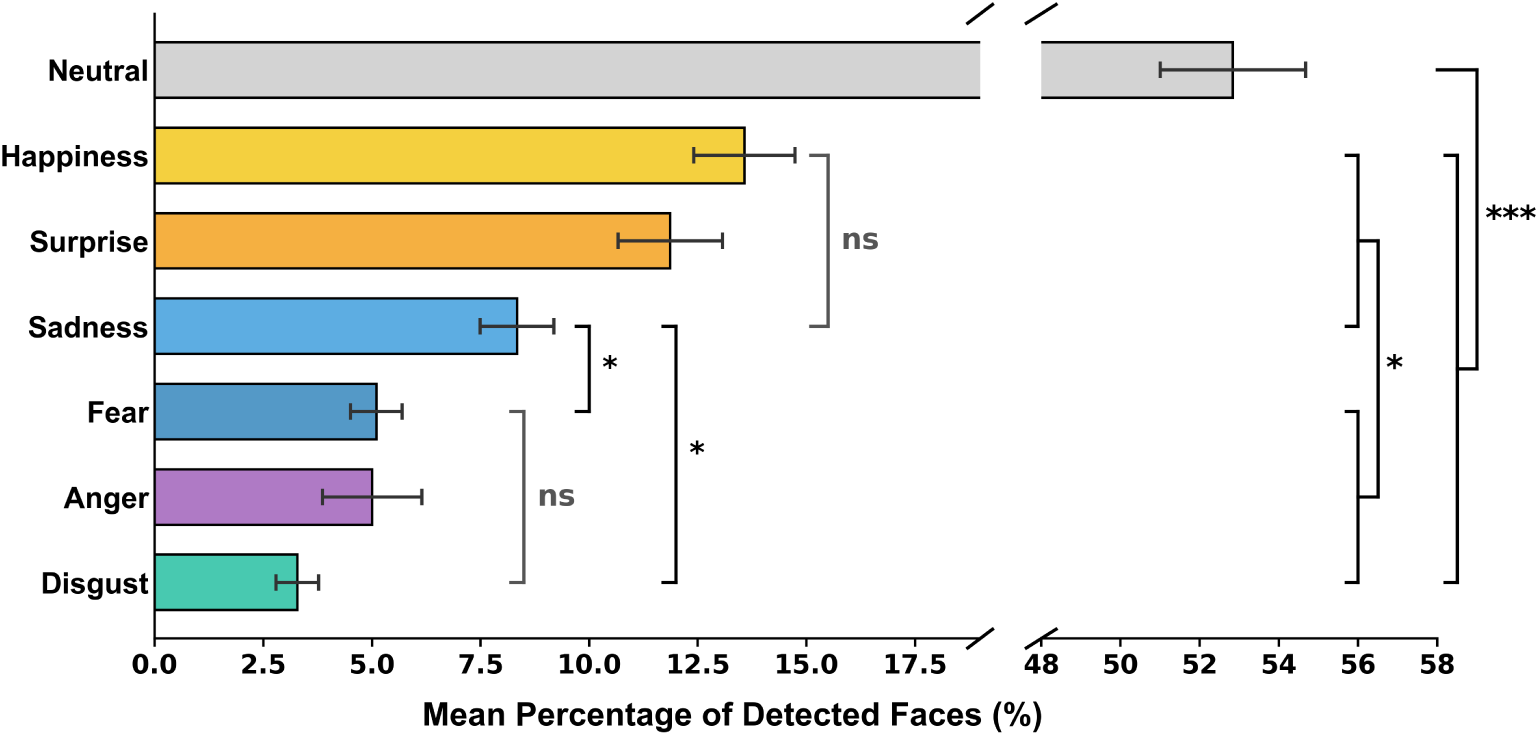
Prevalence of facial emotion expressions in the toddler view (*N*=26). The X-axis is split to accommodate the scale difference between emotional facial expressions and neutral. The error bars represent *±*1 SE. *ns* -indicates pairwise comparisons that are not statistically significant; *** - *α < .*001, * - min *α < .*05 (Bonferroni-corrected).

Post-hoc pairwise comparisons (Bonferroni corrected) revealed three distinct prevalence groups: *Neutral* expressions occurred significantly more frequently than all other categories (*p < .*001); *Happiness* and *Surprise* occurred significantly more frequently than *Anger*, *Fear*, and *Disgust*; and no significant differences occurred between *Anger*, *Fear*, and *Disgust*. Among the negative expressions, *Sadness* occupied an intermediate position, differing significantly from *Disgust* and *Fear*, but not from *Anger* (Figure 5 and Supplementary Table S1).

### 3.3 Presence of Emotional Expressions in the Caregiver View

We next examined the facial emotional expressions present in the caregiver visual input, focusing on the 25 families with valid caregiver footage. Similar to the toddler view, a one-way repeated measures ANOVA confirmed a significant effect of facial expression category, *F*(3.07, 73.56) = 220.97, *p < .*001, *η_p_*^2^ = .902. Emotionally *Neutral* facial expressions were the most prevalent (*M* = 53.97%, *SD* = 9.31%). Among the emotional expressions, *Happiness* was most prevalent (*M* = 19.58%, *SD* = 7.20%), followed by *Surprise* (*M* = 9.47%, *SD* = 5.17%). The negative expressions remained the least frequent categories: *Sadness* (*M* = 7.70%, *SD* = 5.55%); *Anger* (*M* = 5.98%, *SD* = 5.83%); *Fear* (*M* = 2.26%, *SD* = 1.32%) and *Disgust* (*M* = 1.04%, *SD* = 1.07%) (Figure 6).

**Figure 6:**
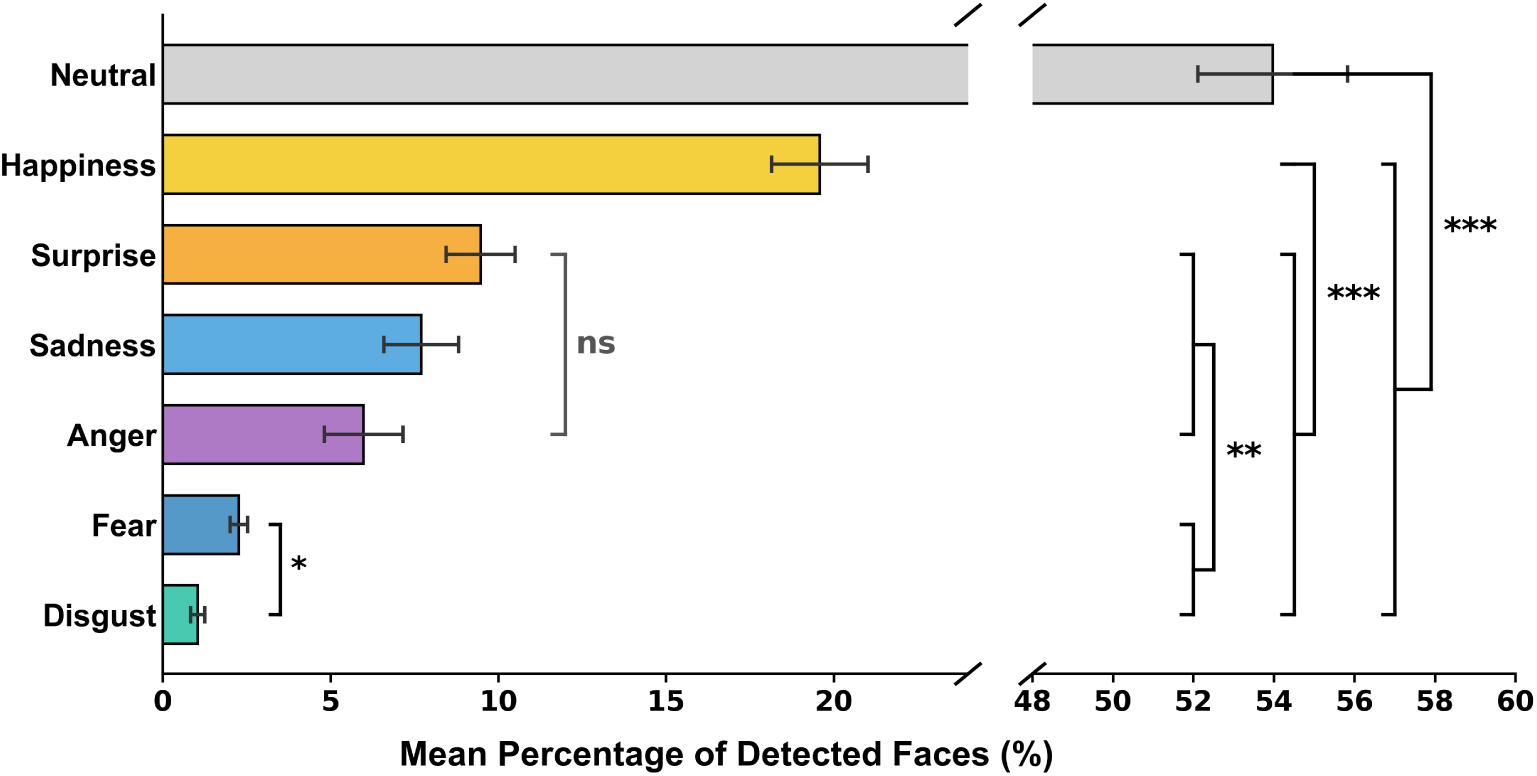
Prevalence of facial emotion expressions in the caregiver view (*N*=25). The X-axis is split to accommodate the scale difference between emotional facial expressions and neutral. The error bars represent *±*1 SE. *ns* - indicates pairwise comparisons that are not statistically significant; *** - *α < .*001, ** - minimum *α < .*01, * - *α < .*05 (Bonferroni-corrected).

Post-hoc pairwise comparisons (Bonferroni-corrected) indicated that *Neutral* expressions occurred significantly more often than all other categories. Furthermore, *Happiness* appeared significantly more frequently than *Surprise* and all negative emotions. Among the negative expressions, *Sadness* and *Anger* were significantly more frequent than *Disgust*, while *Fear* did not differ significantly from *Anger* (Figure 6 and Supplementary Table S2).

### 3.4 Comparing Facial Expression Presence between the Toddler and Caregiver View

Lastly, we directly compared the occurrence of facial emotional expressions between the toddler and caregiver perspectives by using the subsample with concurrent recordings (*N* = 24). For these analyses, we examined facial expressions as percentages of all face-opportunities.

A 2 (Perspective: *Toddler*, *Caregiver*) *×* 7 (Facial Expression: *Anger*, *Disgust*, *Fear*, *Happiness*, *Neutral*, *Sadness*, *Surprise*) repeated-measures ANOVA revealed a significant main effect of expression (*F*(1.36, 31.33) = 75.27, *p < .*001, *η_p_*^2^ = .766) and a significant main effect of perspective (*F*(1.00, 23.00) = 24.10, *p < .*001, *η_p_*^2^ = .512). These main effects were further qualified by a significant interaction, *F*(1.63, 37.50) = 18.21, *p < .*001, *η_p_*^2^ = .442.

Given our aim to study potential differences between perspectives, we calculated pairwise comparisons (Bonferroni-corrected) between the toddler and caregiver view for each expression. The results indicated that the caregiver’s view contained a significantly higher occurrence of *Happiness* (*M* = 2.67, *SE* = .32) compared to the toddler’s view (*M* = 1.04%, *SE* = .13, *p < .*001). A similar pattern was observed for emotionally *Neutral* faces, which were more frequent for caregivers (*M* = 8.18%, *SE* = .94) compared to toddlers (*M* = 4.39%, *SE* = .53, *p < .*001). Crucially, the caregiver’s perspective also captured significantly more expressions of *Sadness* (*M* = 1.15%, *SE* = .21) compared to the toddler’s view (*M* = 0.71%, *SE* = .12, *p* = .042). No statistically significant differences were found for *Anger*, *Disgust*, *Fear*, or *Surprise* (Table 2 and Figure 7).

**Figure 7:**
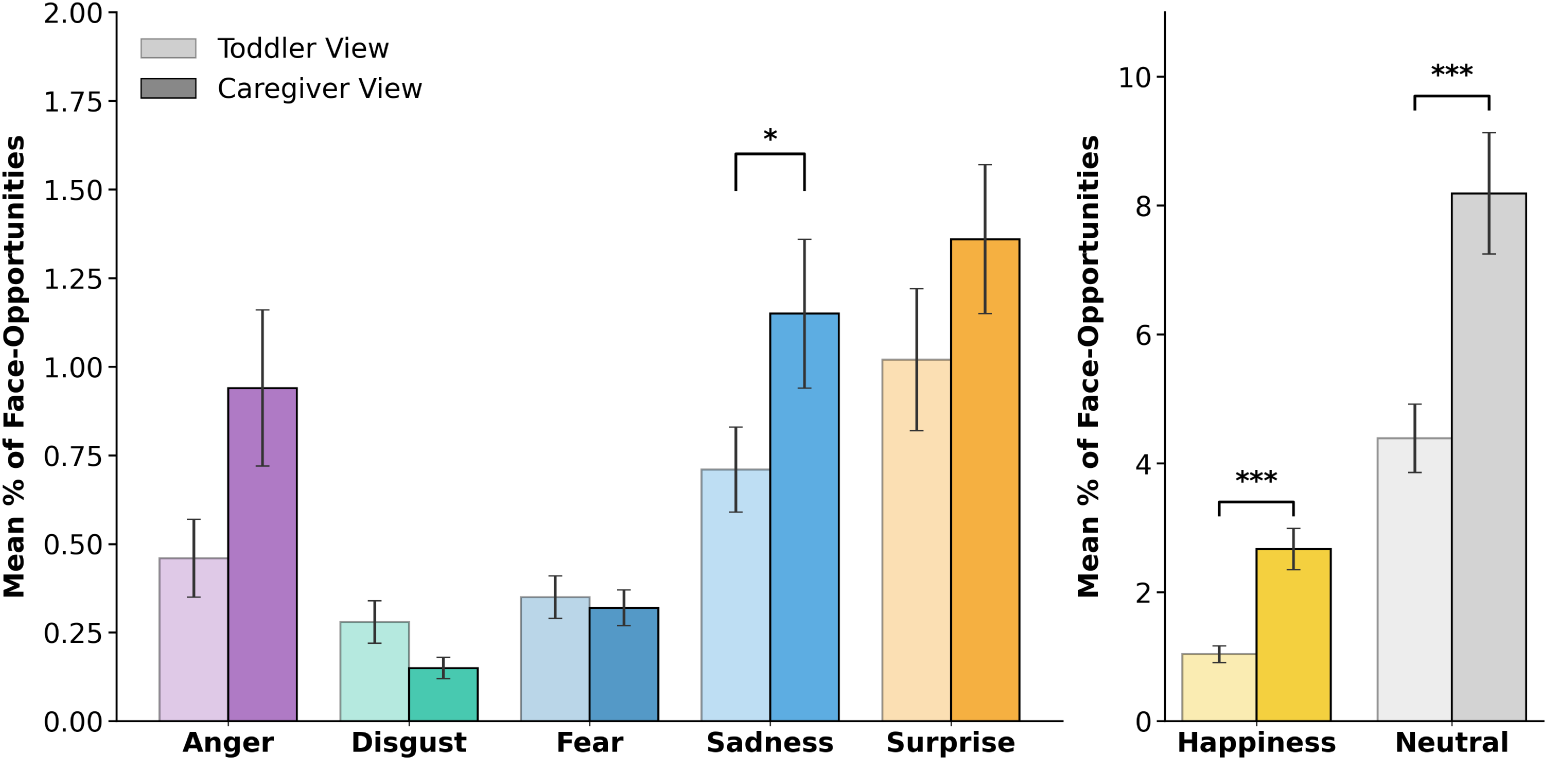
Prevalence of facial emotion expressions in the toddler and caregiver view. The bar chart represents the percentage of frames in the toddler and caregiver views containing faces assigned each facial emotion expression (*N* = 24 families with corresponding toddler and caregiver data). Lighter colors = toddler view; darker colors = caregiver view. For improved readability, *Happiness* and *Neutral* are illustrated in a separate panel with a larger y-axis scale. The error bars represent +*/ −* 1*SE*. * *p < .*05, *** *p < .*001.

**Table 2:** The results of paired-samples comparisons of facial emotion expressions between toddler and caregiver views (Estimates and Pairwise Comparisons).

| Expression | Mean Percentage (SE) |  | Comparisons |  |
| --- | --- | --- | --- | --- |
|  | Toddler View | Caregiver View | *p*-value | Significant? |
| Anger | 0.46 (0.11) | 0.94 (0.22) | .058 | No |
| Disgust | 0.28 (0.06) | 0.15 (0.03) | .062 | No |
| Fear | 0.35 (0.06) | 0.32 (0.05) | .690 | No |
| Sadness | 0.71 (0.12) | 1.15 (0.21) | .042 | Yes |
| Surprise | 1.02 (0.20) | 1.36 (0.21) | .073 | No |
| Happiness | 1.04 (0.13) | 2.67 (0.32) | < .001 | Yes |
| Neutral | 4.39 (0.53) | 8.18 (0.94) | < .001 | Yes |

*Note.* Mean and Standard Error (SE) values are based on estimated marginal means. Significance determined using Bonferroni adjustment for multiple comparisons.

## 4 Discussion

It has been long recognized that the real world visual input is important for developing abilities to process facial expressions of emotion (e.g., Johnson, 2011; Leppänen and Nelson, 2009; Morton and Johnson, 1991). However, thus far, little to no direct evidence existed to characterize the nature of this input and how infants and toddlers learn from it. Leveraging technological advances in wearable sensing for toddlers, and computer vision, the present study provides, to our knowledge, the first objective quantification of the emotional facial expressions toddlers encounter in their daily home environment. By capturing the dual perspectives of the toddler and the caregiver simultaneously, our results reveal both similarities and differences between the two partners. Regarding the frequency of faces, we replicated established findings that faces are relatively sparse in both the toddlers’ egocentric view (approx. 7.5%), and in the view of caregivers (approx. 13.8%) (e.g., Fausey et al., 2016; Jayaraman et al., 2015, 2017; Jayaraman and Smith, 2019; Oruc et al., 2019). Crucially, our analyses reveal that this sparsity extends to the emotional signal itself.

Several previous studies using head-mounted cameras, the majority of which were conducted in the USA and Canada, have reported that toddlers’ egocentric views are characterized by a distinct drop in face prevalence relative to younger infants (e.g., Fausey et al., 2016; Long et al., 2022a). Separately, research with adults suggests that this sparsity of faces may be unique to toddlers and younger children. For example, Oruc et al. (2019) showed that in a variety of contexts, Canadian adults encounter faces at rates similar to those of younger infants, suggesting a non-linear developmental trajectory of face exposure. Critically, we report data for a substantial sample of UK participants, recorded across a variety of events in the home environment. Our study replicates the toddlerhood dip in face frequency, and further demonstrates that the difference between toddlers and adults persists even when controlling for the environment. While our absolute frequency rates are numerically lower than previous reports (likely due to the conservative nature of our automated detection, which was tuned to filter out ambiguous or temporally unstable detections), the relative pattern remains robust: in the home, adults have faces in view at rates comparable to when high-density environments like the workplace are included, and these rates are higher than those of toddlers.

Why does this divergence occur? Given that social opportunities (the number of people in the home) was held constant between toddler and caregiver, the sparsity of faces in the toddler view cannot be attributed to a lack of social partners. Instead, it likely stems from how toddlers interact with the environment. On one hand, this may be motivational. As toddlers transition from face-learning to face-using (e.g., social referencing), prolonged looks may become less necessary. Furthermore, as manual exploration increases, attention is increasingly allocated to hands and objects as primary sources of learning (Yu and Smith, 2013). On the other hand, biomechanical constraints likely play a significant role. While the social environment is shared, the physical vantage point is not. Due to height differences, adults can view faces by looking level or down, a movement supported by gravity and greater ocular range of motion (Lee et al., 2019). In contrast, toddlers must look up to see faces, a motor action that is more energy-costly and biomechanically restricted. Additionally, the resource load of early walking imposes a functional constraint: novice walkers must frequently direct their gaze downward to navigate the path ahead, leaving less opportunity to orient toward faces high in the visual environment (Kretch et al., 2014). Thus, the toddler dip in face exposure may be as much the cascading effect of physical maturity as it is of development in other areas of social and cognitive development.

Critically, for the first time, we show that within this already limited exposure to faces, the affective visual signal is sparse. For both toddlers and caregivers, faces conveying emotional expressions represent less than half of the total number of faces occurring in their egocentric view. Furthermore, negative facial expressions are under-represented, while happiness emerges as the most dominant facial expression. The sparsity of the affective facial signal is even more obvious when considering the wider context of all visual input in the natural home environment. Emotional facial displays appear in toddlers’ view less than 4% of the time, and in the caregiver’s less than 7%. Previous literature, primarily reflecting on empirical data from adult studies, has suggested that in the natural environment, adults seldom spontaneously manifest facial expressions of affect, and these displays tend to be more subtle and fleeting than the exaggerated posed expressions typically used in laboratory experiments (e.g., Crivelli and Fridlund, 2019; Dawel et al., 2023; Cowen and Keltner, 2020; Steward et al., 2025). Consistent with these suggestions, our study provides direct evidence that in the natural home environment, toddlers and caregivers from a Western culture have limited opportunities to observe and process facial displays of affect. Furthermore, the ecological validity of these findings is emphasized by the scale and breadth of our sampling. Unlike studies relying on prescribed play situations or short observation windows, the current dataset captures a range of home events across several hours of observations and a substantial cohort. This suggests that the observed sparsity is not an artifact of short observation windows or specific activity contexts, but a stable feature of the toddlers’ and caregivers’ visual ecology. These findings are critical for reframing our understanding of how affect knowledge develops.

The finding that caregivers encounter significantly more expressions of happiness and sadness than toddlers aligns with research on the functional nature of early emotional signaling. Buss and Kiel (2004) demonstrated that during distressing situations, toddlers display more sadness compared to fear and anger, and they do so specifically when looking at their mother. This may be for the purpose of eliciting support, consistent with other studies which show that parents are more likely to comfort and assist their children when they are expressing sadness (Corapci et al., 2025). Consequently, the caregiver’s first-person view is naturally characterized by an increased frequency of sadness. Furthermore, toddlers are also less likely to down regulate intense positive affect and excitement (e.g., Cole, 1986; Cole et al., 2004). In contrast, the toddler’s view captures the caregiver, who is likely regulating their own expression to appear calm or reassuring and have internalized display rules (e.g., Corapci et al., 2025; Denham et al., 2004; Lin et al., 2021; Malatesta et al., 1986), resulting in an observed asymmetry that reflects the functional regulation of the dyad.

Interestingly, the dominance of happy and surprise, and the sparsity of negative facial displays like fear and disgust, in toddlers’ egocentric view mirrors broad trends in children’s emotion understanding development. For example, in an extensive meta-analysis, including children 2- to 12-years-old, Riddell et al. (2024) have shown that across a variety of laboratory-based tasks and stimuli, the ability to explicitly recognize emotional expressions improves throughout childhood, but, at all ages, happiness was the most easily recognized emotion category, followed by anger, surprise, sadness, disgust, and fear. One possible interpretation, in line with our findings, is that the relative predominance of happy expressions in the visual input throughout infancy and childhood contributes to its robust recognition. In contrast, the extreme sparsity of negative expressions (e.g., disgust and fear) aligns with their delayed acquisition (e.g. Widen and Russell, 2015). This suggests that the developmental timeline of explicit emotion recognition may be significantly constrained by the opportunities to learn that characterize the child’s natural visual input.

However, the development of emotion processing likely relies on complementary mechanisms that go beyond simple cumulative exposure to facial configurations. A wealth of evidence indicates that infants are sensitive to facial expressions of anger and fear early in ontogeny, and that their behavior is modulated by adult expressivity. The classic literature on social referencing, for instance, demonstrates that infants can decode and act upon facial displays of fear or anger to regulate their own exploration in uncertain contexts (e.g., Hoehl et al., 2008; Nava and Turati, 2022; Sorce et al., 1985; Walle et al., 2017). Furthermore, studies have shown that subliminally perceived angry faces elicit increased sympathetic arousal relative to happy faces in infants as young as 3-4-months (e.g. Nava et al., 2016), while from 7-months, infants’ facial responses to others’ fear and anger seem to reflect reliance on the communicative value of these expressions (e.g., Geangu et al., 2016b; Kaiser et al., 2017). Given our findings that negative facial expressions are characterized by statistical sparsity in the home environment, we propose that their representations do not develop through a gradual accumulation of frequent data. Instead, consistent with sparse coding theories (e.g., Olshausen and Field, 1996) and recent proposals for early development, the infant brain likely optimizes efficiency by capitalizing on high-value, salient instances of input (e.g., Amso and Kirkham, 2021; Barrett et al., 2011; Clerkin et al., 2017; Hoemann et al., 2020; Oliva and Torralba, 2007; Smith et al., 2018).

First, while happy faces are reinforced by frequent social reward, negative expressions likely acquire meaning through high-intensity, episodic conditioning. In potential threat situations (e.g., an infant reaching for hot food), the caregiver’s facial expression is often accompanied by immediate, high-arousal cues, such as loud vocalization, a gasp, or swift physical intervention, that elicit a startle or increased arousal in the infant (e.g. Dahl et al., 2014; LoBue et al., 2025; Campos et al., 1989). This surge in physiological arousal may serve as a potent unconditioned stimulus, allowing the infant to rapidly associate the caregivers’ face with the aversive internal state (e.g., Amso and Kirkham, 2021). These learning mechanisms may be supported by an early emerging emotion-processing neural network, including the orbitofrontal cortex, the amygdala, and the posterior superior temporal sulcus (e.g., Leppänen and Nelson, 2009).

Second, these sparse facial signals are most likely embedded within a rich multimodal and contextual information stream. Real-world emotional expressions are rarely unimodal events; they are coupled with distinct vocal prosody, body postures (e.g., Crespo-Llado et al., 2018; Ke et al., 2022; Ruba and Repacholi, 2020; Reschke et al., 2018; Riviere et al., 2026; Vuong and Geangu, 2023), and embedded within dynamic sequences of actions and events (e.g., Addabbo and Turati, 2020; Addabbo et al., 2026; Aviezer et al., 2017; Hepach and Westermann, 2013; Poulin-Dubois et al., 2018; Reschke et al., 2017b; Roberti et al., 2025; Sacheli et al., 2023) that may follow certain statistical regularities. When facial cues are sparse or transient, the visual system may leverage the gist of the scene or the succession of events (e.g., Barrett et al., 2011; Oliva and Torralba, 2007; Steward et al., 2025). Infants are also adept statistical learners (e.g., Aslin et al., 1998; Bulf et al., 2011; Mermier et al., 2022; Nencheva et al., 2025), capable of detecting these cross-modal regularities and using them to guide their processing of facial signals (e.g., Bahrick and Lickliter, 2010; Ruba and Repacholi, 2020). For example, a fearful face may be learned not by seeing it 1,000 times in isolation, but by reliably detecting its co-occurrence with a fearful voice and body posture or a specific dangerous context (e.g., Crespo-Llado et al., 2018; Dahl et al., 2014; Geangu et al., 2016b; Geangu and Vuong, 2020, 2023; LoBue and Ogren, 2022; Vuong and Geangu, 2023). This multimodality likely scaffolds the learning process (e.g., Bahrick and Lickliter, 2010; Bayet et al., 2018; Emberson et al., 2015; Wang et al., 2025). Indeed, Bayet (2022) argues that top-down signals, including verbal labels and social context, play a critical role in organizing the developing visual cortex, supporting the formation of rich emotional representations even when the visual facial input alone is fleeting or rare.

### 4.1 Limitations and future directions

Our findings provide the first quantitative characterization of the facial affect landscape in the toddler’s natural home environment. However, several limitations warrant consideration and point toward critical avenues for future research.

First, while our automated computer vision pipeline allowed the analysis of a large dataset that would be unfeasible to code manually, it is subject to the constraints of current algorithm performance. To ensure reliability, we employed conservative confidence thresholds to filter out ambiguous or temporally unstable detections. Consequently, our reported frequencies likely represent a lower bound of face and emotion exposure. It is highly probable that partial faces and subtle, low-intensity expressions are more frequent than reported here. However, a key advantage of our validating approach is that the extent of this underestimation can be directly quantified relative to the specific characteristics of the egocentric footage. We argue that as the field moves toward automated quantification of natural experience, such systematic validation across diverse data, devices, and environments must become standard (see also Charlot et al., 2025; Long et al., 2022a; Tang et al., 2025).

It is also important to note that both the human naive perceivers and the computational model exhibited specific patterns of confusion between emotion categories (e.g., between anger and disgust; fear and sadness), with implications for how we estimate the extent of exposure. One might argue that these confusions stem from the image quality limitations inherent to low-cost wearable cameras, which could hinder reliable detection. However, the patterns of confusion observed in our study closely align with those reported in prior research using high-fidelity stimuli in controlled laboratory settings (e.g., Jack et al., 2014, 2016; Woodard et al., 2022). This suggests that the automated model is not merely failing due to noise, but is capturing the inherent perceptual ambiguity of certain facial configurations, effectively quantifying the social input through a human-like affective lens.

As part of a larger effort to develop scalable methodologies for studying social development with high ecological validity (e.g., Geangu et al., 2023; Long et al., 2022a), future research could leverage emerging corpora of egocentric audio-video footage to extend the validation dataset we used in this study. Specifically, training models directly on the unique visual statistics of infantperspective data is expected to further improve robustness. This is particularly critical for under-represented categories. For instance, our validation set contained fewer examples of fear, reflecting the well-documented difficulty of capturing spontaneously elicited fearful facial expressions in naturalistic context (see also Dupré et al. (2020)). Furthermore, while discrete classification is necessary for establishing the absolute prevalence of emotion categories (as performed here), future research could complement this by shifting toward using the distributional probabilities of the facial expressive signal. This would allow researchers to specifically investigate the ambiguity of the input, characterizing the subtle, mixed, and context-dependent affective displays from which infants and toddlers are actually learning.

Second, our study focused exclusively on the home environment. While the home is the primary ecological niche in the early years, it is not the only learning environment. It remains to be seen whether the scarcity of faces and negative affect persists in other settings, such as the nursery, playgrounds, or busy public spaces. It is plausible that these environments offer a face-dense counterpart to the home, characterized by different regularities, such as a higher frequency of peer interactions, conflict, and associated negative affect. A comprehensive account of the early visual input requires mapping these varied contexts to understand how the input shapes the development of emotion expression processing.

Third, our sample consisted of families from a Western cultural background (UK). As highlighted in the introduction, cultural norms significantly influence affect display rules and infant information sampling strategies (e.g., Geangu et al., 2016a; Jack et al., 2012). In cultures that prioritize higher physical proximity and different socialization goals, the frequency and nature of emotional signaling may differ substantially. Extending this dual-perspective approach to nonWestern samples is crucial for distinguishing universal characteristics of the visual input from the culturally specific ones.

Finally, a central proposal of our discussion is that complementary mechanisms may be involved in learning from sparse visual emotional signals, and that these rely on multimodal affective signal and physiological salience. However, the present analysis was restricted to the visual modality. We did not quantify the acoustic environment or the caregiver’s vocal prosody, and we did not consider toddlers’ behavioral and physiological responses. Also, we did not analyze the temporal structure of the visual affective signal. As argued in recent frameworks for a temporal science of behaviour (e.g., Abney et al., 2025), developmental inputs often follow non-linear, bursty dynamics, as seen in the auditory domain (e.g., Warlaumont et al., 2022), which may be critical for learning. Testing hypotheses derived from this proposal requires integrating visual with audio processing, and synchronized infant physiological responses. Novel platforms of integrated wearable sensors, such as EgoActive and LittleBeats (e.g., Geangu et al., 2023; Islam et al., 2024; Mason et al., 2024; Zhang et al., 2024) and recent advances in voice-speech technologies (e.g., Laudańska et al., 2025; Charlot et al., 2025; Tang et al., 2025) are promising solutions for capturing this complexity.

Fully exploiting rich naturalistic datasets will require analytic frameworks that go beyond standard frequency metrics. Methodologies such as Representational Similarity Analysis (RSA) (e.g., Diedrichsen and Kriegeskorte, 2017) and dimensionality reduction techniques like Uniform Manifold Approximation and Projection (UMAP) (e.g., McInnes et al., 2018) offer powerful means to map the statistical structure of the input to the structure of the child’s developing representations, testing how the input and the developing mind mutually shape each other. However, capturing the true dynamic nature of development requires moving beyond static analyses to multidimensional modeling of the learning process itself. Future research can potentially model learning and development at multiple time scales, from seconds, to hours, days, and months. In this context, advanced machine learning paradigms, such as continual learning (e.g., Carreira et al., 2024; Lesort et al., 2020; Holton et al., 2025; Parisi et al., 2019; Zeng et al., 2019), become essential. Unlike static models, such learning algorithms could simulate how infants incrementally develop and refine emotion representations over months of exposure, adapting to the changing statistics of their environment. These modeling techniques could allow us to test precisely how the specific sparsity and salience of the input documented here interact with the child’s own physiological arousal to drive learning and development over time.

### 4.2 Conclusions

In conclusion, the present study reveals for the first time that the egocentric view of toddlers and their caregivers is characterized by sparsity rather than abundance of facial emotion expressions. These findings emphasize the need to re-evaluate the mechanisms driving early development: rather than relying on frequent inputs, emotion information processing development likely depends on the efficient exploitation of rare, high-value signals, supported by the broader communicative context. Our study joins a growing body of work demonstrating that moving the study of development from laboratory simulations to the natural environment gives us unprecedented opportunities to anchor our theories of social development in the concrete realities of the child’s lived experience.

## Acknowledgements

We thank all families who participated in this study. We would also like to thank William AP Smith for his technical support. This work was supported by the Wellcome Trust Institutional Strategic Support Fund (ISSF) and the Engineering and Physical Sciences Research Council (EPSRC) Impact Acceleration Account awarded to the University of York (allocated to Elena Geangu), and by the Wellcome Leap 1kD Program (awarded to Elena Geangu).

## Supplementary Information

### A. Supplementary Methods

#### A.0.1. Pipeline implementation

##### Face detection and Emotion Expression Estimation

Faces were detected independently in each frame *𝐼_𝑡_*. For each frame, RetinaFace (ResNet-50) performs a single-stage, multi-task prediction, where it outputs three coupled variables for every candidate face: (1) Bounding Box (*𝑏*): four coordinates (*𝑥*_min_*, 𝑦*_min_*, 𝑥*_max_*, 𝑦*_max_) defining the face region; (2) Confidence Score (*𝑠*): a scalar probability indicating the likelihood of a face; and (3) Facial Landmarks (*𝑙*): a set of five coordinate pairs (*𝑥, 𝑦*) corresponding to the optical centers of the left/right eyes, nose tip, and left/right mouth corners. Next, the landmarks (*𝑙*) were used to compute a similarity transformation matrix, warping the face onto a standardized 112 × 112 pixel reference frame, correcting for in-plane rotation (roll) and scale variations. This step ensures that the subsequent identity and emotion models received spatially normalized inputs.

The aligned RGB tensors (112 × 112 × 3) served as input for two parallel networks. The ResNet-50 architecture extracted a high-dimensional embedding vector to represent facial identity. The EfficientNet-B0 architecture estimated facial expression. We replaced the model’s final classification head with a fully connected layer that outputs a raw logit vector *𝑧* ∈ ℝ^7^. A Softmax activation function was applied to generate a normalized probability distribution P across the seven categories, such that ∑*_i=1_*^7^ 𝑝*_𝑖_* = 1. All raw features (coordinates, embeddings, and emotion probabilities) were archived at this stage prior to filtering.

##### Temporal Check and Quality Filter

To ensure analysis was performed only on valid social cues, the archived detections underwent a two-stage filtering process. A static threshold removed any detections with a bounding box confidence score *<* 0.7. The custom 1D CNN analyzes the time series of confidence scores (*𝑠*) to filter false positives over a 10-frame window. Detections were retained only when meeting a dual-validation criterion: spatial confidence *𝑠 >* 0.7 and temporal probability *𝑝*_CNN_ *>* 0.5.

##### Face Tracking and Temporal Smoothing

In the next step, filtered detections were grouped into short-term clusters (tracks) by optimizing a pairwise affinity matrix. This method calculates affinity using a combination of spatial overlap (Intersection over Union) and identity embedding similarity, a deep feature vector representing the unique biometric appearance of the face. The clustering process was governed by two strict logic gates (constraints) to prevent identity mixing: (1) *Must-not-link constraints*, strictly applied to simultaneous detections within the same frame index *𝑡*. Since a single individual cannot physically appear twice in the same image, any two detections co-occurring in time were prohibited from belonging to the same cluster; and (2) *Must-link constraints*, applied to detections in distinct frames that exhibited high identity embedding similarity and plausible spatial proximity. The algorithm generated up to 20 candidate face tracks. We note that this tracking module was optimized for short-term temporal smoothing rather than long-term re-identification. While identity switches may occur over extended durations, this approach ensured the downstream emotion analysis was performed on temporally continuous face segments, minimizing frame-level noise.

Finally, the emotion probability distributions were temporally smoothed (10-frame moving average) within each track. This approach preserves the model’s uncertainty estimates and mitigates frame-level noise before final classification. This was further used to establish a categorical classification as explained in the next sections.

### B. Supplementary Tables

**Table S1.** Summary of Significant Pairwise Comparisons for Toddler Facial Expressions.

| Comparison | Mean Diff. | SE | p-value | 95% CI |
| --- | --- | --- | --- | --- |
| *Comparisons with Neutral* |  |  |  |  |
| Neutral vs. Happiness | 39.26 | 2.40 | < .001 | [31.15, 47.38] |
| Neutral vs. Surprise | 40.98 | 2.72 | < .001 | [31.79, 50.16] |
| Neutral vs. Sadness | 44.50 | 2.18 | < .001 | [37.15, 51.85] |
| Neutral vs. Fear | 47.74 | 2.17 | < .001 | [40.39, 55.08] |
| Neutral vs. Anger | 47.83 | 2.32 | < .001 | [40.00, 55.67] |
| Neutral vs. Disgust | 49.56 | 2.07 | < .001 | [42.55, 56.57] |
| *Comparisons with Happiness* |  |  |  |  |
| Happiness vs. Anger | 8.57 | 1.91 | .003 | [2.12, 15.02] |
| Happiness vs. Disgust | 10.30 | 1.24 | < .001 | [6.10, 14.49] |
| Happiness vs. Fear | 8.48 | 1.30 | < .001 | [4.09, 12.87] |
| *Comparisons with Surprise* |  |  |  |  |
| Surprise vs. Anger | 6.86 | 1.87 | .024 | [0.55, 13.17] |
| Surprise vs. Disgust | 8.58 | 1.20 | < .001 | [4.53, 12.64] |
| Surprise vs. Fear | 6.76 | 1.26 | < .001 | [2.50, 11.03] |
| *Comparisons with Sadness* |  |  |  |  |
| Sadness vs. Disgust | 5.06 | 1.07 | .002 | [1.44, 8.68] |
| Sadness vs. Fear | 3.24 | 0.88 | .023 | [0.28, 6.20] |

Note. Mean Difference is measured in percentage. SE = Standard Error.

**Table S2.** Summary of Significant Pairwise Comparisons for Caregiver Facial Expressions.

| Comparison | Mean Diff. | SE | p-value | 95% CI |
| --- | --- | --- | --- | --- |
| *Comparisons with Neutral* |  |  |  |  |
| Neutral vs. Disgust | 52.93 | 1.97 | < .001 | [46.24, 59.61] |
| Neutral vs. Fear | 51.72 | 1.85 | < .001 | [45.43, 58.01] |
| Neutral vs. Anger | 47.99 | 2.50 | < .001 | [39.50, 56.48] |
| Neutral vs. Sadness | 46.27 | 2.54 | < .001 | [37.65, 54.89] |
| Neutral vs. Surprise | 44.51 | 2.40 | < .001 | [36.34, 52.67] |
| Neutral vs. Happiness | 34.39 | 2.79 | < .001 | [24.91, 43.88] |
| *Comparisons with Happiness* |  |  |  |  |
| Happiness vs. Disgust | 18.53 | 1.49 | < .001 | [13.47, 23.60] |
| Happiness vs. Fear | 17.32 | 1.52 | < .001 | [12.15, 22.49] |
| Happiness vs. Anger | 13.60 | 2.22 | < .001 | [6.04, 21.15] |
| Happiness vs. Sadness | 11.88 | 1.89 | < .001 | [5.47, 18.28] |
| Happiness vs. Surprise | 10.11 | 1.72 | < .001 | [4.26, 15.96] |
| *Comparisons with Surprise* |  |  |  |  |
| Surprise vs. Disgust | 8.42 | 1.09 | < .001 | [4.74, 12.11] |
| Surprise vs. Fear | 7.21 | 1.10 | < .001 | [3.49, 10.93] |
| *Comparisons with Sadness* |  |  |  |  |
| Sadness vs. Disgust | 6.66 | 1.08 | < .001 | [2.98, 10.33] |
| Sadness vs. Fear | 5.44 | 1.19 | .003 | [1.39, 9.50] |
| *Comparisons with Anger* |  |  |  |  |
| Anger vs. Disgust | 4.94 | 1.04 | .002 | [1.40, 8.47] |
| *Comparisons with Fear* |  |  |  |  |
| Fear vs. Disgust | 1.21 | 0.32 | .017 | [0.13, 2.29] |

Note. Mean Difference is measured in percentage of faces. SE = Standard Error. CI = Confidence Interval. Significance determined using Bonferroni adjustment for multiple comparisons.

